# Power laws and critical fragmentation in global forests

**DOI:** 10.1101/091751

**Authors:** Leonardo A. Saravia, Santiago R. Doyle, Ben Bond-Lamberty

## Abstract

The replacement of forest areas with human-dominated landscapes usually leads to fragmentation, altering the structure and function of the forest. Here we studied the dynamics of forest patch sizes at a global level, examining signals of a critical transition from an unfragmented to a fragmented state, using the MODIS vegetation continuous field. We defined wide regions of connected forest across continents and big islands, and combined five criteria, including the distribution of patch sizes and the fluctuations of the largest patch over the last sixteen years, to evaluate the closeness of each region to a fragmentation threshold. Regions with the highest deforestation rates—South America, Southeast Asia, Africa—all met these criteria and may thus be near a critical fragmentation threshold. This implies that if current forest loss rates are maintained, wide continental areas could suddenly fragment, triggering extensive species loss and degradation of ecosystems services.

## Introduction

Forests are one of the most important biomes on earth, providing habitat for a large proportion of species and contributing extensively to global biodiversity (Crowther *et al*., 2015). In the previous century, human activities have influenced global bio-geochemical cycles (Bonan, 2008; Canfield *et al*., 2010), with one of the most dramatic changes being the replacement of 40% of Earth’s formerly biodiverse land areas with landscapes that contain only a few species of crop plants, domestic animals and humans (Foley *et al*., 2011). These local changes have accumulated over time and now constitute a global forcing (Barnosky *et al*., 2012). Another global scale forcing that is tied to habitat destruction is fragmentation, which is defined as the division of a continuous habitat into separated portions that are smaller and more isolated. Fragmentation produces multiple interwoven effects: reductions of biodiversity between 13% and 75%, decreasing forest biomass, and changes in nutrient cycling (Haddad *et al*., 2015). The effects of fragmentation are not only important from an ecological point of view but also that of human activities, as ecosystem services are deeply influenced by the level of landscape fragmentation (Rudel *et al*., 2005; Angelsen, 2010; Mitchell *et al*., 2015).

Ecosystems have complex interactions between species and present feedbacks at different levels of organization (Gilman *et al*., 2010), and external forcings can produce abrupt changes from one state to another, called critical transitions (Scheffer *et al*., 2009). Such ‘critical’ transitions have been detected mostly at local scales (Drake & Griffen, 2010; Carpenter *et al*., 2011), but the accumulation of changes in local communities that overlap geographically can propagate and theoretically cause an abrupt change of the entire system at larger scales (Barnosky *et al*., 2012).

Complex systems can experience two general classes of critical transitions (Solé, 2011). In so-called first-order transitions, a catastrophic regime shift that is mostly irreversible occurs because of the existence of alternative stable states (Scheffer *et al*., 2001). This class of transitions is suspected to be present in a variety of ecosystems such as lakes, woodlands, coral reefs (Scheffer *et al*., 2001), semi-arid grasslands (Bestelmeyer *et al*., 2011), and fish populations (Vasilakopoulos & Marshall, 2015). They can be the result of positive feedback mechanisms (Villa Martín *et al*., 2015); for example, fires in some forest ecosystems were more likely to occur in previously burned areas than in unburned places (Kitzberger *et al*., 2012).

The other class of critical transitions are continuous or second order transitions (Solé & Bascompte, 2006). In these cases, there is a narrow region where the system suddenly changes from one domain to another, with the change being continuous and in theory reversible. This kind of transitions were suggested to be present in tropical forests (Pueyo *et al*., 2010; Taubert *et al*., 2018), semi-arid mountain ecosystems (McKenzie & Kennedy, 2012), and tundra shrublands (Naito & Cairns, 2015). The transition happens at a critical point where we can observe a distinctive spatial pattern: scale-invariant fractal structures characterized by power law patch distributions (Stauffer & Aharony, 1994). There are several processes that can convert a catastrophic transition to a second order transition (Villa Martín *et al*., 2015). These include stochasticity, such as demographic fluctuations, spatial heterogeneities, and/or dispersal limitation. All these components are present in forest around the globe (Seidler & Plotkin, 2006; Filotas *et al*., 2014; Fung *et al*., 2016), and thus continuous transitions might be more probable than catastrophic transitions. Moreover, there is some evidence of recovery in some systems that supposedly suffered an irreversible transition produced by overgrazing (Zhang *et al*., 2005; Bestelmeyer *et al*., 2013) and desertification (Allington & Valone, 2010).

The spatial phenomena observed in continuous critical transitions deal with connectivity, a fundamental property of general systems and ecosystems from forests (Ochoa-Quintero *et al*., 2015) to marine ecosystems (Leibold & Norberg, 2004) and the whole biosphere (Lenton & Williams, 2013). When a system goes from a fragmented to a connected state we say that it percolates (Solé, 2011). Percolation implies that there is a path of connections that involve the whole system. Thus we can characterize two domains or phases: one dominated by short-range interactions where information cannot spread, and another in which long-range interactions are possible and information can spread over the whole area. (The term “information” is used in a broad sense and can represent species dispersal or movement.) Thus, there is a critical “percolation threshold” between the two phases, and the system could be driven close to or beyond this point by an external force; climate change and deforestation are the main forces that could be the drivers of such a phase change in contemporary forests (Bonan, 2008; Haddad *et al*., 2015). There are several applications of this concept in ecology: species’ dispersal strategies are influenced by percolation thresholds in three-dimensional forest structure (Solé *et al*., 2005), and it has been shown that species distributions also have percolation thresholds (He & Hubbell, 2003). This implies that pushing the system below the percolation threshold could produce a biodiversity collapse (Bascompte & Solé, 1996; Solé *et al*., 2004; Pardini *et al*., 2010); conversely, being in a connected state (above the threshold) could accelerate the invasion of the forest into prairie (Loehle *et al*., 1996; Naito & Cairns, 2015).

One of the main challenges with systems that can experience critical transitions—of any kind—is that the value of the critical threshold is not known in advance. In addition, because near the critical point a small change can precipitate a state shift in the system, they are difficult to predict. Several methods have been developed to detect if a system is close to the critical point, e.g. a deceleration in recovery from perturbations, or an increase in variance in the spatial or temporal pattern (Hastings & Wysham, 2010; Carpenter *et al*., 2011; Boettiger & Hastings, 2012; Dai *et al*., 2012).

The existence of a critical transition between two states has been established for forest at a global scale in different works (Hirota *et al*. (2011); Staal *et al*. (2016); Wuyts *et al*. (2017)). It is generally believed that this constitutes a first order catastrophic transition. The regions where forest can grow are not distributed homogeneously, as there are demographic fluctuations in forest growth and disturbances produced by human activities. All these factors imply that if these were first order transitions they will be converted or observed as second order continuous transitions (Villa Martín *et al*., 2014, 2015). From this basis, we applied indices derived from second-order transitions to global forest cover dynamics.

In this study, our objective is to look for evidence that forests around the globe are near continuous critical points that represent a fragmentation threshold. We use the framework of percolation to first evaluate if forest patch distribution at a continental scale is described by a power law distribution and then examine the fluctuations of the largest patch. The advantage of using data at a continental scale is that for very large systems the transitions are very sharp (Solé, 2011) and thus much easier to detect than at smaller scales, where noise can mask the signals of the transition.

## Methods

### Study areas definition

We analyzed mainland forests at a continental scale, covering the whole globe, by delimiting land areas with a near-contiguous forest cover, separated with each other by large non-forested areas. Using this criterion, we delimited the following forest regions. In America, three regions were defined: South America temperate forest (SAT), subtropical and tropical forest up to Mexico (SAST), and USA and Canada forest (NA). Europe and North Asia were treated as one region (EUAS). The rest of the delimited regions were South-east Asia (SEAS), Africa (AF), and Australia (OC). We also included in the analysis islands larger than 10^5^km^2^. We applied this criterion to delimit regions because we based our study on percolation theory that assumes some kind of connectivity in the study area (Appendix Table S2, figure S1-S6).

### Forest patch distribution

We studied forest patch distribution in each defined area from 2000 to 2015 using the MODerate-resolution Imaging Spectroradiometer (MODIS) Vegetation Continuous Fields (VCF) Tree Cover dataset version 051 (DiMiceli *et al*., 2015). This dataset is produced at a global level with a 231-m resolution, from 2000 onwards on an annual basis, the last available year was 2015. There are several definitions of forest based on percent tree cover (Sexton *et al*., 2015); we choose a range from 20% to 40% threshold in 5% increments to convert the percentage tree cover to a binary image of forest and non-forest pixels. This range is centered in the definition used by the United Nations’ International Geosphere-Biosphere Programme (Belward, 1996), and studies of global fragmentation (Haddad *et al*., 2015) and includes the range used in other studies of critical transitions (Xu *et al*., 2016). Using this range we try to avoid the errors produced by low discrimination of MODIS VCF between forest and dense herbaceous vegetation at low forest cover and the saturation of MODIS VCF in dense forests (Sexton *et al*., 2013). We repeat all the analysis for this set of thresholds, except in some specific cases described below. Patches of contiguous forest were determined in the binary image by grouping connected forest pixels using a neighborhood of 8 forest units (Moore neighborhood). The MODIS VCF product defines the percentage of tree cover by pixel but does not discriminate the type of trees so besides natural forest it includes plantations of tree crops like rubber, oil palm, eucalyptus and other managed stands (Hansen *et al*., 2014).

### Percolation theory

A more in-depth introduction to percolation theory can be found elsewhere (Stauffer & Aharony, 1994) and a review from an ecological point of view is available (Oborny *et al*., 2007). Here, to explain the basic elements of percolation theory we formulate a simple model: we represent our area of interest by a square lattice and each site of the lattice can be occupied—e.g. by forest—with a probability *p*. The lattice will be more occupied when *p* is greater, but the sites are randomly distributed. We are interested in the connection between sites, so we define a neighborhood as the eight adjacent sites surrounding any particular site. The sites that are neighbors of other occupied sites define a patch. When there is a patch that connects the lattice from opposite sides, it is said that the system percolates. When *p* is increased from low values, a percolating patch suddenly appears at some value of *p* called the critical point *p*_*c*_.

Thus percolation is characterized by two well-defined phases: the unconnected phase when *p < p*_*c*_ (called subcritical in physics), in which species cannot travel far inside the forest, as it is fragmented; in a general sense, information cannot spread. The second is the connected phase when *p > p*_*c*_ (supercritical), species can move inside a forest patch from side to side of the area (lattice), i.e. information can spread over the whole area. Near the critical point, several scaling laws arise: the structure of the patch that spans the area is fractal, the size distribution of the patches is power-law, and other quantities also follow power-law scaling (Stauffer & Aharony, 1994).

The value of the critical point *p*_*c*_ depends on the geometry of the lattice and on the definition of the neighborhood, but other power-law exponents only depend on the lattice dimension. Close to the critical point, the distribution of patch sizes is:

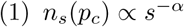

where *n*_*s*_(*p*) is the number of patches of size *s*. The exponent *α* does not depend on the details of the model and it is called universal (Stauffer & Aharony, 1994). These scaling laws can be applied to landscape structures that are approximately random, or at least only correlated over short distances (Gastner *et al*., 2009). In physics, this is called “isotropic percolation universality class”, and corresponds to an exponent *α* = 2.05495. If we observe that the patch size distribution has another exponent it will not belong to this universality class and some other mechanism should be invoked to explain it. Percolation thresholds can also be generated by models that have some kind of memory (Hinrichsen, 2000; Ódor, 2004): for example, a patch that has been exploited for many years will recover differently than a recently deforested forest patch. In this case, the system could belong to a different universality class, or in some cases, there is no universality, in which case the value of *α* will depend on the parameters and details of the model (Corrado *et al*., 2014).

To illustrate these concepts, we conducted simulations with a simple forest model with only two states: forest and non-forest. This type of model is called a “contact process” and was introduced for epidemics (Harris, 1974) but has many applications in ecology (Solé & Bascompte, 2006; Gastner *et al*., 2009). A site with forest can become extinct with probability *e*, and produce another forest site in a neighborhood with probability *c*. We use a neighborhood defined by an isotropic power law probability distribution. We defined a single control parameter as *λ*= *c/e* and ran simulations for the subcritical fragmentation state *λ* < *λ*_c_, with *λ*= 2, near the critical point for *λ*= 2.5, and for the supercritical state with *λ*= 5 (see supplementary data, gif animations).

### Patch size distributions

We fitted the empirical distribution of forest patches calculated for each of the thresholds on the range we previously mentioned. We used maximum likelihood (Goldstein *et al*., 2004; Clauset *et al*., 2009) to fit four distributions: power-law, power-law with exponential cut-off, log-normal, and exponential. We assumed that the patch size distribution is a continuous variable that was discretized by the remote sensing data acquisition procedure.

The power-law distribution needs a lower bound for its scaling behavior. This lower bound is also estimated from the data by maximizing the Kolmogorov-Smirnov (KS) statistic, computed by comparing the empirical and fitted cumulative distribution functions (Clauset *et al*., 2009). For the log-normal model, we constrain the values of the *µ* parameter to positive values, this parameter controls the mode of the distribution and when is negative most of the probability density of the distribution lies outside the range of the forest patch size data (Limpert *et al*., 2001).

To select the best model we calculated corrected Akaike Information Criteria (*AIC*_*c*_) and Akaike weights for each model (Burnham & Anderson, 2002). Akaike weights (*w*_*i*_) are the weight of evidence in favor of model *i* being the actual best model given that one of the *N* models must be the best model for that set of *N* models. Additionally, we computed a likelihood ratio test (Vuong, 1989; Clauset *et al*., 2009) of the power law model against the other distributions. We calculated bootstrapped 95% confidence intervals (Crawley, 2012) for the parameters of the best model, using the bias-corrected and accelerated (BCa) bootstrap (Efron & Tibshirani, 1994) with 10000 replications.

### Largest patch dynamics

The largest patch is the one that connects the highest number of sites in the area. This has been used extensively to indicate fragmentation (Gardner & Urban, 2007; Ochoa-Quintero *et al*., 2015) and the size of the largest patch *S*_*max*_ has been studied in relation to percolation phenomena (Stauffer & Aharony, 1994; Bazant, 2000; Botet & Ploszajczak, 2004) but is seldom used in ecological studies (for an exception see Gastner *et al*. (2009)). When the system is in a connected state (*p > p*_*c*_) the landscape is almost insensitive to the loss of a small fraction of forest, but close to the critical point a minor loss can have important effects (Solé & Bascompte, 2006; Oborny *et al*., 2007), because at this point the largest patch will have a filamentary structure, i.e. extended forest areas will be connected by thin threads. Small losses can thus produce large fluctuations.

One way to evaluate the fragmentation of the forest is to calculate the proportion of the largest patch against the total area (Keitt *et al*., 1997). The total area of the regions we are considering (Appendix S4, figures S1-S6) may not be the same as the total area that the forest could potentially occupy, and thus a more accurate way to evaluate the weight of *S*_*max*_ is to use the total forest area, which can be easily calculated by summing all the forest pixels. We calculate the proportion of the largest patch for each year, dividing *S*_*max*_ by the total forest area of the same year: 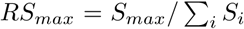. This has the effect of reducing the *S*_*max*_ fluctuations produced due to environmental or climatic changes influences in total forest area. When the proportion *RS*_*max*_ is large (more than 60%) the largest patch contains most of the forest so there are fewer small forest patches and the system is probably in a connected phase. Conversely, when it is low (less than 20%), the system is probably in a fragmented phase (Saravia & Momo, 2018). To define if a region will be in a connected or unconnected state we used the *RS*_*max*_ of the highest (i.e., most conservative) threshold of 40%, that represents the densest area of forest within our chosen range. We assume that there are two alternative states for the critical transition—the forest could be fragmented or unfragmented. If *RS*_*max*_ is a good indicator of the fragmentation state of the forest its distribution of frequencies should be bimodal (Bestelmeyer *et al*., 2011), so we apply the Hartigan’s dip test that measures departures from unimodality (Hartigan & Hartigan, 1985).

To evaluate if the system is near a critical transition, we calculate the fluctuations of the largest patch *∆S*_*max*_ = *S*_*max*_(*t*) − 〈*S*_*max*_〉, using the same formula for *RS*_*max*_. To characterize fluctuations we fitted three empirical distributions: power-law, log-normal, and exponential, using the same methods described previously. We expect that large fluctuations near a critical point have heavy tails (log-normal or power-law) and that fluctuations far from a critical point have exponential tails, corresponding to Gaussian processes (Rooij *et al*., 2013). We also apply likelihood ratio test explained previously (Vuong, 1989; Clauset *et al*., 2009); if the p-values obtained to compare the best distribution against the others are not significant we concluded that there is not enough data to decide which is the best model. We generated animated maps showing the fluctuations of the two largest patches at 30% threshold, to aid in the interpretations of the results.

A robust way to detect if the system is near a critical transition is to analyze the increase in variance of the density (Benedetti-Cecchi *et al*., 2015)—in our case ‘density’ is the total forest cover divided by the area. It has been demonstrated that the variance increase in density appears when the system is very close to the transition (Corrado *et al*., 2014), and thus practically it does not constitute an early warning indicator. An alternative is to analyze the variance of the fluctuations of the largest patch ∆*S*_*max*_: the maximum is attained at the critical point but a significant increase occurs well before the system reaches the critical point (Corrado *et al*., 2014; Saravia & Momo, 2018). In addition, before the critical fragmentation, the skewness of the distribution of ∆*S*_*max*_ should be negative, implying that fluctuations below the average are more frequent. We characterized the increase in the variance using quantile regression: if variance is increasing the slopes of upper or/and lower quartiles should be positive or negative.

All statistical analyses were performed using the GNU R version 3.3.0 (R Core Team, 2015), to fit the distributions of patch sizes we used the Python package powerlaw (Alstott *et al*., 2014). For the quantile regressions, we used the R package quantreg (Koenker, 2016). Image processing was done in MATLAB r2015b (The Mathworks Inc.). The complete source code for image processing and statistical analysis and the patch size data files are available at figshare http://dx.doi.org/10.6084/m9.figshare.4263905.

## Results

Figure 1 shows an example of the distribution of the biggest 200 patches for years 2000 and 2014. This distribution is highly variable; the biggest patch usually maintains its spatial location, but sometimes it breaks and then big temporal fluctuations in its size are observed, as we will analyze below. Smaller patches can merge or break more easily so they enter or leave the list of 200, and this is why there is a color change across years.

**Figure 1:**
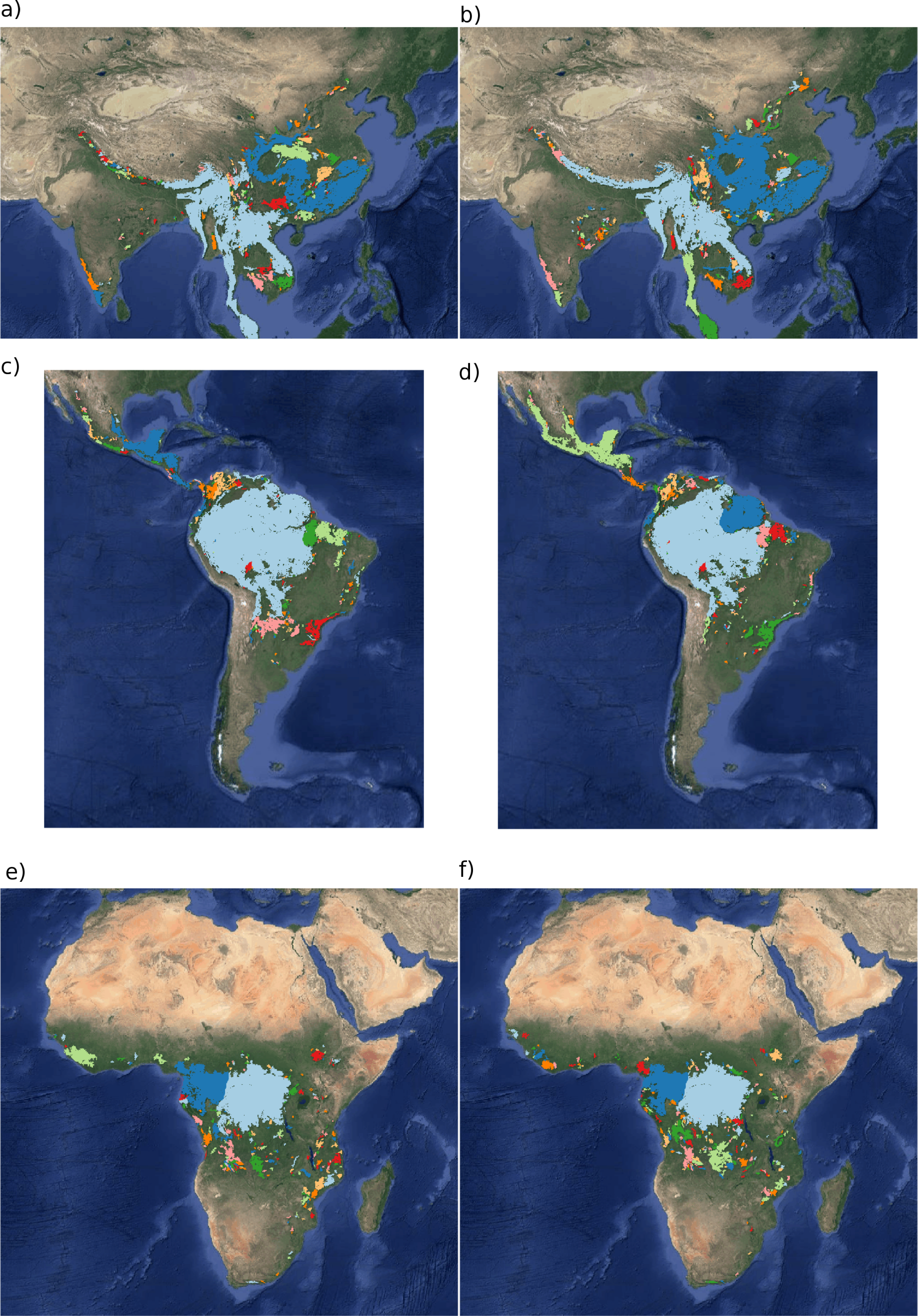
Forest patch distributions for continental regions for the years 2000 and 2014. The images are the 200 biggest patches, shown at a coarse pixel scale of 2.5 km. The regions are: a) & b) southeast Asia; c) & d) South America subtropical and tropical and e) & f) Africa mainland, for the years 2000 and 2014 respectively. The color palette was chosen to discriminate different patches and does not represent patch size.

**Figure 2:**
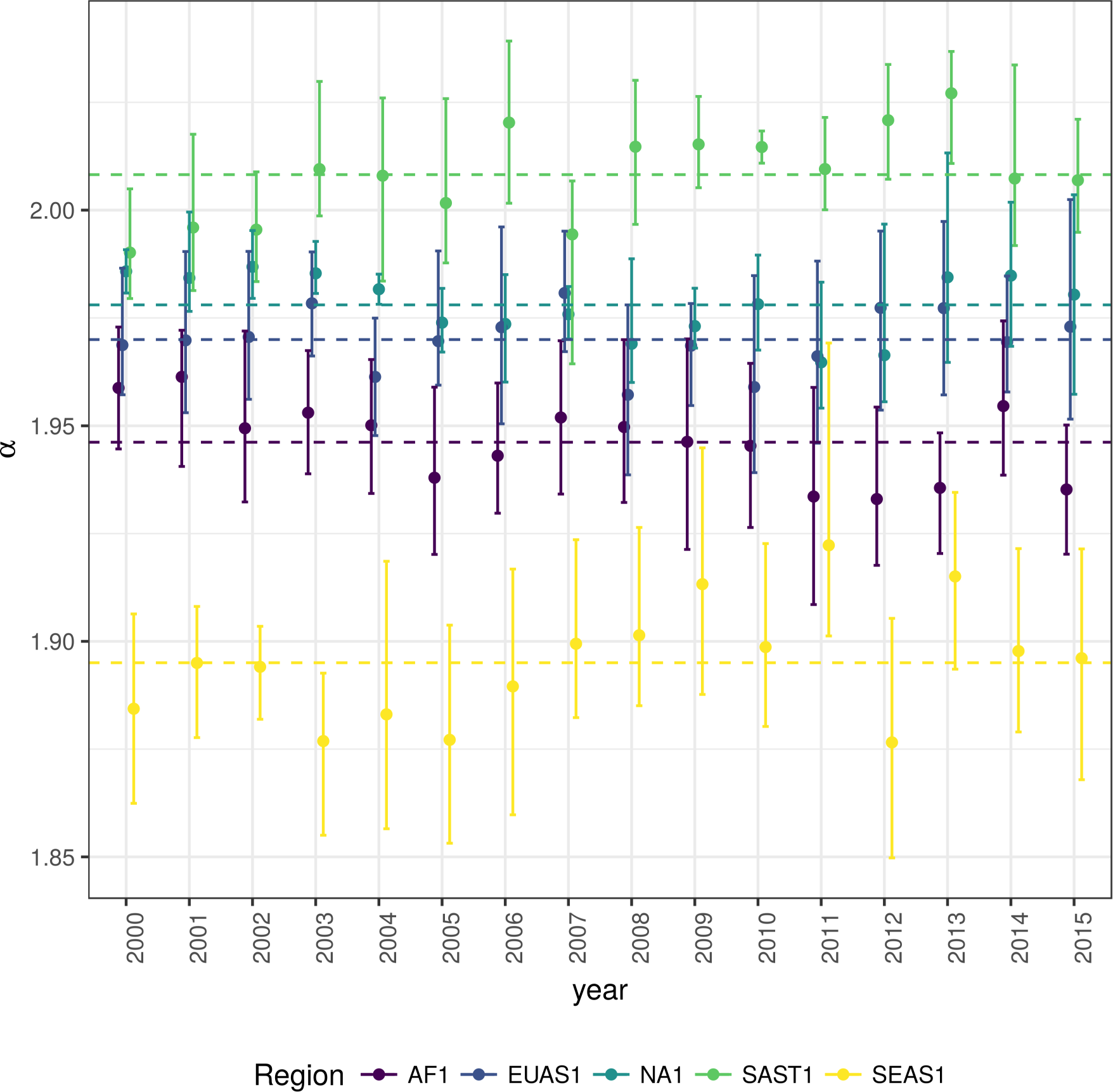
Power law exponents (*α*) of forest patch distributions for regions with total forest area *>* 10^7^ km^2^. Dashed horizontal lines are the means by region, with 95% confidence interval error bars estimated by bootstrap resampling. The regions are AF1: Africa mainland, EUAS1: Eurasia mainland, NA1: North America mainland, SAST1: South America subtropical and tropical, SEAS1: Southeast Asia mainland.

The power law distribution was selected as the best model in 99% of the cases (Figure S7). In a small number of cases (1%), the power law with exponential cutoff was selected, but the value of the parameter *α* was similar by ±0.03 to the pure power law (Table S1 and model fit data table). Additionally, the patch size where the exponential tail begins is very large, and thus we used the power law parameters for these cases (region EUAS3, SAST2). In finite-size systems, the favored model should be the power law with exponential cut-off because the power-law tails are truncated to the size of the system (Stauffer & Aharony, 1994). This implies that differences between the two kinds of power law models should be small. We observe this effect: when the pure power-law model is selected as the best model the likelihood ratio test shows that in 64% of the cases the differences with power law with exponential cutoff are not significant (p-value>0.05); in these cases the differences between the fitted *α* for both models are less than 0.001. Instead, the likelihood ratio test clearly differentiates the power law model from the exponential model (100% cases p-value<0.05), and the log-normal model (90% cases p-value<0.05).

The global mean of the power-law exponent *α* is 1.967 and the bootstrapped 95% confidence interval is 1.964 - 1.970. The global values for each threshold are different, because their confidence intervals do not overlap, and their range goes from 1.90 to 2.01 (Table S1). Analyzing the biggest regions (Figure 1, Table S2) the northern hemisphere regions (EUAS1 & NA1) have similar values of *α* (1.97, 1.98), pantropical areas have different *α* with greatest values for South America (SAST1, 2.01) and in descending order Africa (AF1, 1.946) and Southeast Asia (SEAS1, 1.895). With greater *α* the fluctuations of patch sizes are lower and vice versa (Newman, 2005).

We calculated the total areas of forest and the largest patch *S*_*max*_ by year for different thresholds, and as expected these two values increase for smaller thresholds (Table S3). We expect fewer variations in the largest patch relative to total forest area *RS*_*max*_ (Figure S9); in ten cases it stayed near or higher than 60% (EUAS2, NA5, OC2, OC3, OC4, OC5, OC6, OC8, SAST1, SAT1) over the 25-35 range or more. In four cases it stayed around 40% or less, at least over the 25-30% range (AF1, EUAS3, OC1, SAST2), and in six cases there is a crossover from more than 60% to around 40% or less (AF2, EUAS1, NA1, OC7, SEAS1, SEAS2). This confirms the criteria of using the most conservative threshold value of 40% to interpret *RS*_*max*_ with regard to the fragmentation state of the forest. The frequency of *RS*_*max*_ showed bimodality (Figure S10) and the dip test rejected unimodality (D = 0.0416, p-value = 0.0003), which also implies that *RS*_*max*_ is a good index to study the fragmentation state of the forest.

The *RS*_*max*_ for regions with more than 10^7^ km^2^ of forest is shown in figure 3. South America tropical and subtropical (SAST1) is the only region with an average close to 60%, the other regions are below 30%. Eurasia mainland (EUAS1) has the lowest value near 20%. For regions with less total forest area (Figure S10, Table S3), Great Britain (EUAS3) has the lowest proportion less than 5%, Java (OC7) and Cuba (SAST2) are under 25%, while other regions such as New Guinea (OC2), Malaysia/Kalimantan (OC3), Sumatra (OC4), Sulawesi (OC5) and South New Zealand (OC6) have a very high proportion (75% or more). Philippines (SEAS2) seems to be a very interesting case because it seems to be under 30% until the year 2007, fluctuates around 30% in years 2008-2010, then jumps near 60% in 2011-2013 and then falls again to 30%, this seems an example of a transition from a fragmented state to an unfragmented one (figure S11).

**Figure 3:**
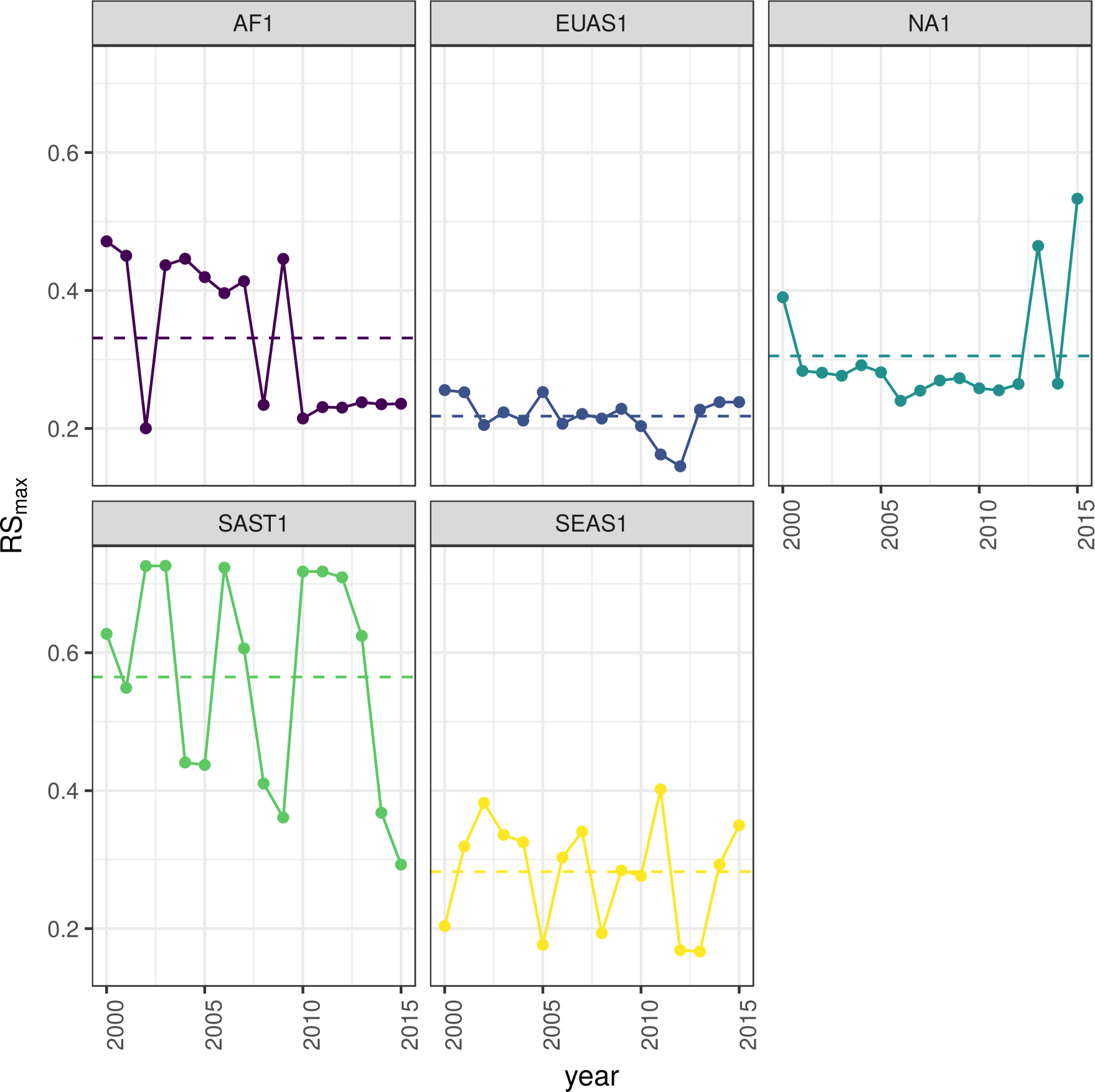
Largest patch proportion relative to total forest area *RS*_*max*_, for regions with total forest area *>*10^7^ km^2^. We show here the *RS*_*max*_ calculated using a threshold of 40% of forest in each pixel to determine patches. Dashed lines are averages across time. The regions are AF1: Africa mainland, EUAS1: Eurasia mainland, NA1: North America mainland, SAST1: South America tropical and subtropical, SEAS1: Southeast Asia mainland.

We analyzed the distributions of fluctuations of the largest patch relative to total forest area ∆*RS*_*max*_ and the fluctuations of the largest patch ∆*S*_*max*_. Although the Akaike criteria identified different distributions as the best, in most cases the Likelihood ratio test was not significant and we are not able, from these data, to determine with confidence which is the best distribution. In only one case was the distribution selected by the Akaike criteria confirmed as the correct model for relative and absolute fluctuations (Table S4).

The animations of the two largest patches (see supplementary data, largest patch gif animations) qualitatively shows the nature of fluctuations and if the state of the forest is connected or not. If the largest patch is always the same patch over time, the forest is probably not fragmented; this happens for regions with *RS*_*max*_ of more than 40% such as AF2 (Madagascar), EUAS2 (Japan), NA5 (Newfoundland) and OC3 (Malaysia). In regions with *RS*_*max*_ between 40% and 30%, the identity of the largest patch could change or stay the same in time. For OC7 (Java) the largest patch changes and for AF1 (Africa mainland) it stays the same. Only for EUAS1 (Eurasia mainland), we observed that the two largest patches are always the same, implying that these two large patches are probably by a geographical accident but they have the same dynamics. The regions with *RS*_*max*_ less than 25% (SAST2-Cuba and EUAS3-Great Britain) have an always-changing largest patch reflecting their fragmented state. In the case of SEAS2 (Philippines), a transition is observed, with the identity of the largest patch first variable, and then constant after 2010.

The results of quantile regressions are almost identical for ∆*RS*_*max*_ and ∆*S*_*max*_ (table S5). Among the biggest regions, Africa (AF1) has a similar pattern across thresholds but only the 30% threshold is significant; the upper and lower quantiles have significant negative slopes, but the lower quantile slope is lower, implying that negative fluctuations and variance are increasing (Figure 4). Eurasia mainland (EUAS1) has significant slopes at 20%, 30% and 40% thresholds but the patterns are different at 20% variance is decreasing, at 30% and 40% only is increasing. Thus the variation of the densest portion of the largest patch is increasing within a limited range. North America mainland (NA1) exhibits the same pattern at 20%, 25% and 30% thresholds: a significant lower quantile with positive slope, implying decreasing variance. South America tropical and subtropical (SAST1) have significant lower quantile with a negative slope at 25% and 30% thresholds indicating an increase in variance. SEAS1 has an upper quantile with positive slope significant for 25% threshold, also indicating an increasing variance. The other regions, with forest area smaller than 10^7^km^2^ are shown in figure S11 and table S5. Philippines (SEAS2) is an interesting case: the slopes of lower quantiles are positive for thresholds 20% and 25%, and the upper quantile slopes are positive for thresholds 30% and 40%; thus variance is decreasing at 20%-25% and increasing at 30%-40%.

**Figure 4:**
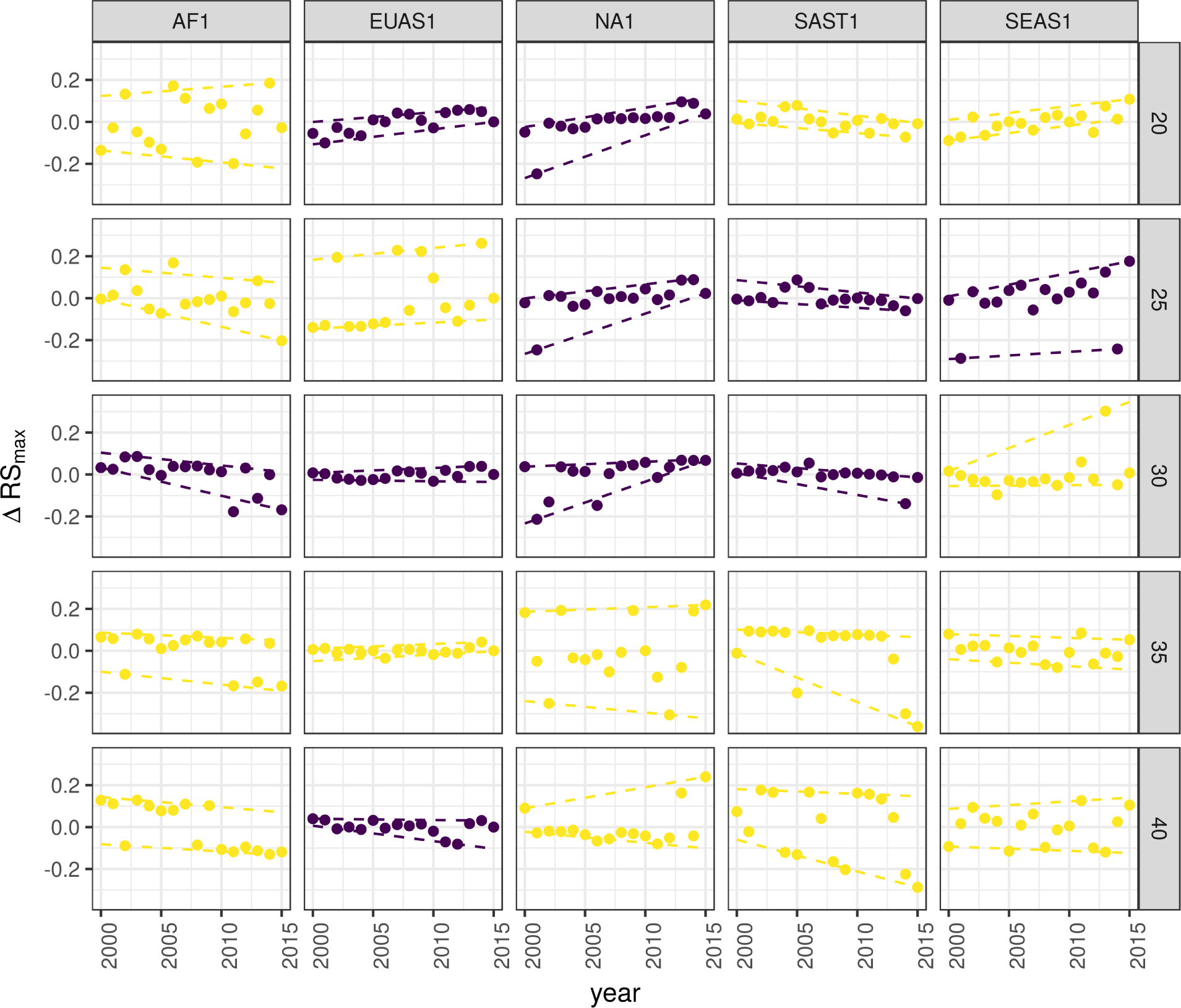
Largest patch fluctuations for regions with total forest area >10^7^km^2^ across years. The patch sizes are relative to the total forest area of the same year. Dashed lines are 90% and 10% quantile regressions, to show if fluctuations were increasing; purple (dark) panels have significant slopes. The regions are AF1: Africa mainland, EUAS1: Eurasia mainland, NA1: North America mainland, SAST1: South America tropical and subtropical, SEAS1: Southeast Asia mainland.

The conditions that indicate that a region is near a critical fragmentation threshold are that patch size distributions follow a power law; variance of ∆*RS*_*max*_ is increasing in time, and skewness is negative. All these conditions must happen at the same time at least for one threshold. When the threshold is higher more dense regions of the forest are at risk. This happens for Africa mainland (AF1), Eurasia mainland(EUAS1), Japan (EUAS2), Australia mainland (OC1), Malaysia/Kalimantan (OC3), Sumatra (OC4), South America tropical & subtropical (SAST1), Cuba (SAST2), Southeast Asia, Mainland (SEAS1).

## Discussion

We found that the forest patch distribution of all regions of the world, spanning tropical rainforests, boreal and temperate forests, followed power laws spanning seven orders of magnitude. Power laws have previously been found for several kinds of vegetation, but never at global scales as in this study. Moreover, the range of the estimated power law exponents is relatively narrow (1.90 - 2.01), even though we tested a variety of different thresholds levels. This suggests the existence of one unifying mechanism, or perhaps different mechanisms that act in the same way in different regions, affecting forest spatial structure and dynamics.

Several mechanisms have been proposed for the emergence of power laws in forests. The first is related self organized criticality (SOC), when the system is driven by its internal dynamics to a critical state; this has been suggested mainly for fire-driven forests (Zinck & Grimm, 2009; Hantson *et al*., 2015). Real ecosystems do not seem to meet the requirements of SOC dynamics (Pueyo *et al*., 2010; McKenzie & Kennedy, 2012), however, because they have both endogenous and exogenous controls, are non-homogeneous, and do not have a separation of time scales (Solé *et al*., 2002; Solé & Bascompte, 2006). A second possible mechanism, suggested by Pueyo et al. (2010), is isotropic percolation: when a system is near the critical point, the power law structures arise. This is equivalent to the random forest model that we explained previously, and requires the tuning of an external environmental condition to carry the system to this point. We did not expect forest growth to be a random process at local scales, but it is possible that combinations of factors cancel out to produce seemingly random forest dynamics at large scales. This has been suggested as a mechanism for the observed power laws of global tropical forest at year 2000 (Taubert *et al*., 2018). In this case we should have observed power laws in a limited set of situations that coincide with a critical point, but instead we observed pervasive power law distributions. Thus isotropic percolation does not seem likely to be the mechanism that produces the observed distributions.

A third possible mechanism is facilitation (Manor & Shnerb, 2008; Irvine *et al*., 2016): a patch surrounded by forest will have a smaller probability of being deforested or degraded than an isolated patch. The model of Scanlon et al. (2007) showed an *α* = 1.34 which is different from our results (1.90 - 2.01 range). Another model but with three states (tree/non-tree/degraded), including local facilitation and grazing, was also used to obtain power laws patch distributions without external tuning, and exhibited deviations from power laws at high grazing pressures (Kéfi *et al*., 2007). The values of the power law exponent *α* obtained for this model are dependent on the intensity of facilitation: when facilitation is more intense the exponent is higher, but the maximal values they obtained are still lower than the ones we observed. Thus an exploration of the parameters of this model and simulations at a continental scale will be needed to find if this is a plausible mechanism.

It has been suggested that a combination of spatial and temporal indicators could more reliably detect critical transitions (Kéfi *et al*., 2014). In this study, we combined five criteria to evaluate the closeness of the system to a fragmentation threshold. Two of them were spatial: the forest patch size distribution, and the proportion of the largest patch relative to total forest area *RS*_*max*_. The other three were the distribution of temporal fluctuations in the largest patch size, the trend in the variance, and the skewness of the fluctuations. One of them: the distribution of temporal fluctuations ∆*RS*_*max*_ can not be applied with our temporal resolution due to the difficulties of fitting and comparing heavy tailed distributions. The combination of the remaining four gives us an increased degree of confidence about the system being close to a critical transition.

Monitoring the biggest patches using *RS*_*max*_ is also important regardless of the existence or not of critical transitions. *RS*_*max*_ is relative to total forest area thus it could be used to compare regions with a different extension of forests and as the total area of forest also changes with different environmental conditions, e.g. this could be used to compare forest from different latitudes. Moreover, the areas covered by *S*_*max*_ across regions contain most of the intact forest landscapes defined by Potapov *et al*. (2008a), and thus *RS*_*max*_ is a relatively simple way to evaluate the risk in these areas.

South America tropical and subtropical (SAST1), Southeast Asia mainland (SEAS1) and Africa mainland (AF1) met all criteria at least for one threshold; these regions generally experience the biggest rates of deforestation with a significant increase in loss of forest (Hansen *et al*., 2013). From our point of view the most critical region is Southeast Asia, because the proportion of the largest patch relative to total forest area *RS*_*max*_ was 28%. This suggests that Southeast Asia’s forests are the most fragmented, and thus its flora and fauna the most endangered. Due to the criteria we adopted to define regions, we could not detect the effect of conservation policies applied at a country level, e.g. the Natural Forest Conservation Program in China, which has produced an 1.6% increase in forest cover and net primary productivity over the last 20 years (Viña *et al*., 2016). Indonesia and Malaysia (OC3) both are countries with hight deforestation rates (Hansen *et al*., 2013); Sumatra (OC4) is the biggest island of Indonesia and where most deforestation occurs. Both regions show a high *RS*_*max*_ greater than 60%, and thus the forest is in an unfragmented state, but they met all other criteria, meaning that they are approaching a transition if the actual deforestation rates continue.

The Eurasian mainland region (EUAS1) is an extensive area with mainly temperate and boreal forest, and a combination of forest loss due to fire (Potapov *et al*., 2008b) and forestry. The biggest country is Russia that experienced the biggest rate of forest loss of all countries, but here in the zone of coniferous forest the the largest gain is observed due to agricultural abandonment (Prishchepov *et al*., 2013). The loss is maximum at the most dense areas of forest (Hansen *et al*., 2013, Table S3), this coincides with our analysis that detect an increasing risk at denser forest. This region also has a relatively low *RS*_*max*_ that means is probably near a fragmented state. A region that is similar in forest composition to EAUS1 is North America (NA1); the two main countries involved, United States and Canada, have forest dynamics mainly influenced by fire and forestry, with both regions are extensively managed for industrial wood production. North America has a higher *RS*_*max*_ than Eurasia and a positive skewness that excludes it from being near a critical transition. A possible explanation of this is that in Russia after the collapse of the Soviet Union harvest was lower due to agricultural abandonment but illegal overharvesting of high valued stands has increased in recent decades (Gauthier *et al*., 2015).

The analysis of *RS*_*max*_ reveals that the island of Philippines (SEAS2) seems to be an example of a critical transition from an unconnected to a connected state, i.e. from a state with high fluctuations and low *RS*_*max*_ to a state with low fluctuations and high *RS*_*max*_. If we observe this pattern backwards in time, the decrease in variance become an increase, and negative skewness is constant, and thus the region exhibits the criteria of a critical transition (Table 1, Figure S12). The actual pattern of transition to an unfragmented state could be the result of an active intervention of the government promoting conservation and rehabilitation of protected areas, ban of logging old-growth forest, reforestation of barren areas, community-based forestry activities, and sustainable forest management in the country’s production forest (Lasco *et al*., 2008). This confirms that the early warning indicators proposed here work in the correct direction. An important caveat is that the MODIS dataset does not detect if native forest is replaced by agroindustrial tree plantations like oil palms, that are among the main drivers of deforestation in this area (Malhi *et al*., 2014). To improve the estimation of forest patches, data sets as the MODIS cropland probability and others about land use, protected areas, forest type, should be incorporated (Hansen *et al*., 2014; Sexton *et al*., 2015).

**Table 1.**
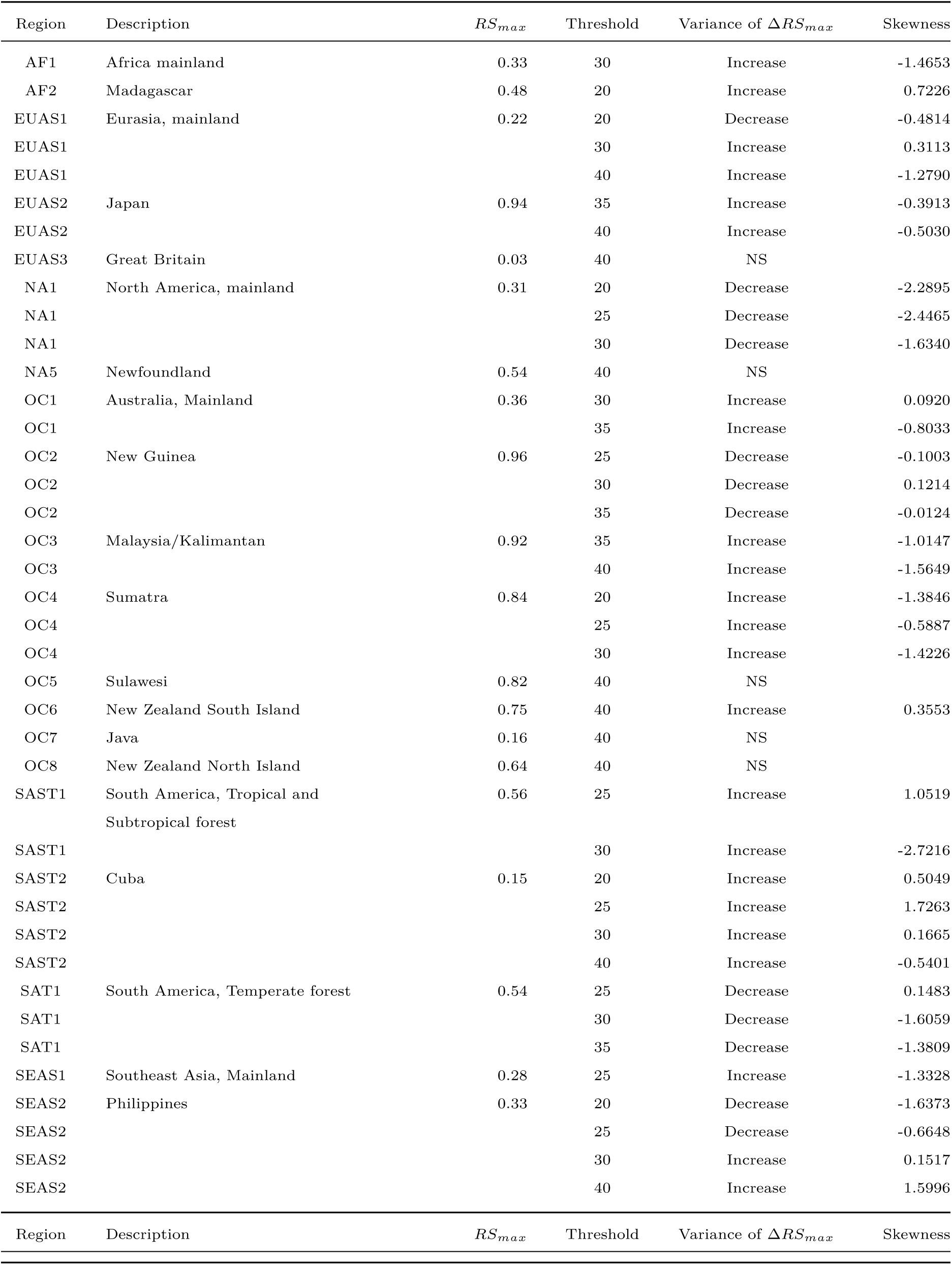
Regions and indicators of closeness to a critical fragmentation point. Where: *RS*_*max*_ is the largest patch divided by the total forest area; Threshold is the value used to calculate patches from the MODIS VCF pixels; ∆*RS*_*max*_ are the fluctuations of *RS*_*max*_ around the mean and the increase or decrease in the variance was estimated using quantile regressions; skewness was calculated for *RS*_*max*_. NS means the results were non-significant. The conditions that determine the closeness to a fragmentation point are: increasing variance of ∆*RS*_*max*_ and negative skewness. *RS*_*max*_ indicates if the forest is unfragmented (>0.6) or fragmented (<0.3).

Deforestation and fragmentation are closely related. At low levels of habitat reduction species population will decline proportionally; this can happen even when the habitat fragments retain connectivity. As habitat reduction continues, the critical threshold is approached and connectivity will have large fluctuations (Brook *et al*., 2013). This could trigger several negative synergistic effects: population fluctuations and the possibility of extinctions will rise, increasing patch isolation and decreasing connectivity (Brook *et al*., 2013). This positive feedback mechanism will be enhanced when the fragmentation threshold is reached, resulting in the loss of most habitat specialist species at a landscape scale (Pardini *et al*., 2010). Some authors have argued that since species have heterogeneous responses to habitat loss and fragmentation, and biotic dispersal is limited, the importance of thresholds is restricted to local scales or even that its existence is questionable (Brook *et al*., 2013). Fragmentation is by definition a local process that at some point produces emergent phenomena over the entire landscape, even if the area considered is infinite (Oborny *et al*., 2005). If a forest is already in a fragmented state, a second critical transition from forest to non-forest could happen: the desertification transition (Corrado *et al*., 2014). Considering the actual trends of habitat loss, and studying the dynamics of non-forest patches—instead of the forest patches as we did here—the risk of this kind of transition could be estimated. The simple models proposed previously could also be used to estimate if these thresholds are likely to be continuous and reversible or discontinuous and often irreversible (Weissmann & Shnerb, 2016), and the degree of protection (e.g. using the set-asides strategy Banks-Leite *et al*. (2014)) that would be necessary to stop this trend.

The effectiveness of landscape management is related to the degree of fragmentation, and the criteria to direct reforestation efforts could be focused on regions near a transition (Oborny *et al*., 2007). Regions that are in an unconnected state require large efforts to recover a connected state, but regions that are near a transition could be easily pushed to a connected state; feedbacks due to facilitation mechanisms might help to maintain this state. Crossing the fragmentation critical point in forests could have negative effects on biodiversity and ecosystem services (Haddad *et al*., 2015), but it could also produce feedback loops at different levels of the biological hierarchy. This means that a critical transition produced at a continental scale could have effects at the level of communities, food webs, populations, phenotypes and genotypes (Barnosky *et al*., 2012). All these effects interact with climate change, thus there is a potential production of cascading effects that could lead to an abrupt climate change with potentially large ecological and economic impact (Alley *et al*., 2003).

Therefore, even if critical thresholds are reached only in some forest regions at a continental scale, a cascading effect with global consequences could still be produced (Reyer *et al*., 2015). The risk of such event will be higher if the dynamics of separate continental regions are coupled (Lenton & Williams, 2013). At least three of the regions defined here are considered tipping elements of the earth climate system that could be triggered during this century (Lenton *et al*., 2008). These were defined as policy relevant tipping elements so that political decisions could determine whether the critical value is reached or not. Thus the criteria proposed here could be used as a more sensitive system to evaluate the closeness of a tipping point at a continental scale. Further improvements will produce quantitative predictions about the temporal horizon where these critical transitions could produce significant changes in the studied systems.

## Supporting information

### Appendix

*Table S1*: Mean power-law exponent and Bootstrapped 95% confidence intervals by threshold.

*Table S2*: Mean power-law exponent and Bootstrapped 95% confidence intervals across thresholds by region and year.

*Table S3*: Mean total patch area; largest patch *S*_*max*_ in km^2^; largest patch relative to total patch area *R*S_*max*_ and 95% bootstrapped confidence interval of *RS*_*max*_, by region and thresholds, averaged across years

*Table S4*: Model selection for distributions of fluctuation of largest patch ∆*S*_*max*_ and largest patch relative to total forest area ∆*RS*_*max*_.

*Table S5*: Quantil regressions of the fluctuations of the largest patch vs year, for 10% and 90% quantils at different pixel thresholds.

*Table S6*: Unbiased estimation of Skewness of fluctuations of the largest patch ∆*S*_*max*_ and fluctuations relative to total forest area ∆*RS*_*max*_.

*Figure S1*: Regions for Africa: Mainland (AF1), Madagascar (AF2).

*Figure S2*: Regions for Eurasia: Mainland (EUAS1), Japan (EUAS2), Great Britain (EUAS3).

*Figure S3*: Regions for North America: Mainland (NA1), Newfoundland (NA5).

*Figure S4*: Regions for Australia and islands: Australia mainland (OC1), New Guinea (OC2), Malaysia/Kalimantan (OC3), Sumatra (OC4), Sulawesi (OC5), New Zealand south island (OC6), Java (OC7), New Zealand north island (OC8).

*Figure S5*: Regions for South America: Tropical and subtropical forest up to Mexico (SAST1), Cuba (SAST2), South America Temperate forest (SAT1).

*Figure S6*: Regions for Southeast Asia: Mainland (SEAS1), Philippines (SEAS2).

*Figure S7*: Proportion of best models selected for patch size distributions using the Akaike criterion.

*Figure S8*: Power law exponents for forest patch distributions by year for all regions.

*Figure S9*: Average largest patch relative to total forest area *RS*_*max*_ by threshold, for all regions.

*Figure s10*: Frequency distribution of Largest patch proportion relative to total forest area *RS*_*max*_ calculated using a threshold of 40%.

*Figure S11*: Largest patch relative to total forest area *RS*_*max*_ by year at 40% threshold, for regions with total forest area less than 10^7^ km^2^.

*Figure S12*: Fluctuations of largest patch relative to total forest area *RS*_*max*_ for regions with total forest area less than 10^7^ km^2^ by year and threshold.

## Data Accessibility

The MODIS VCF product is freely available from NASA at https://search.earthdata.nasa.gov/. Csv text file with model fits for patch size distribution, and model selection for all the regions; Gif Animations of a forest model percolation; Gif animations of largest patches; patch size files for all years and regions used here; and all the R, Python and Matlab scripts are available at figshare http://dx.doi.org/10.6084/m9.figshare.4263905.

## Acknowledgments

LAS and SRD are grateful to the National University of General Sarmiento for financial support. We want to thank to Jordi Bascompte, Nara Guisoni, Fernando Momo, and two anonymous reviewers for their comments and discussions that greatly improved the manuscript. This work was partially supported by a grant from CONICET (PIO 144-20140100035-CO).

